# HLA3DB: comprehensive annotation of peptide/HLA complexes enables blind structure prediction of T cell epitopes

**DOI:** 10.1101/2023.03.20.533510

**Authors:** Sagar Gupta, Santrupti Nerli, Sreeja Kutti Kandy, Glenn L. Mersky, Nikolaos G. Sgourakis

## Abstract

The class I proteins of the major histocompatibility complex (MHC-I) display epitopic peptides derived from endogenous proteins on the cell surface for immune surveillance. Accurate modeling of peptide/HLA (pHLA, the human MHC) structures has been mired by conformational diversity of the central peptide residues, which are critical for recognition by T cell receptors. Here, analysis of X-ray crystal structures within a curated database (HLA3DB) shows that pHLA complexes encompassing multiple HLA allotypes present a discrete set of peptide backbone conformations. Leveraging these representative backbones, we employ a regression model trained on terms of a physically relevant energy function to develop a comparative modeling approach for nonamer peptide/HLA structures named RepPred. Our method outperforms the top pHLA modeling approach by up to 19% in terms of structural accuracy, and consistently predicts blind targets not included in our training set. Insights from our work provide a framework for linking conformational diversity with antigen immunogenicity and receptor cross-reactivity.

## Main

The class I major histocompatibility complex (MHC-I) protein presents self, foreign, or aberrant peptides to provide a mechanism of immune surveillance via CD8^+^ cytotoxic T cell lymphocytes^1^. The human version of MHC-I, Human Leukocyte Antigen class I (HLA-I), is encoded by the highly polymorphic HLA gene locus^2^. There are three classical (HLA-A, HLA-B, and HLA-C) as well as three non-classical HLA genes (HLA-E, HLA-F, and HLA-G) that combine to encode over 20,000 HLA allotypes^3, 4^. Each allotype can display a repertoire of up to 10^8^ epitopic peptide sequences, defining distinct peptide/HLA-I (pHLA) complexes^5^. The large sequence variation and subsequent differences in peptide repertoires between HLA allotypes can cause disease susceptibility^6^. Additionally, this diversity across allotypes ensures species viability as all antigens can likely be presented by at least one allotype in the population. At the structural level, peptides of length 8 to 15 amino acids bind to the HLA groove which consists of six major pockets labeled from A to F^7^. Canonical binding occurs via stable anchoring interactions within the B and F pockets of the HLA and the second (P2) and last (PΩ) peptide residues, respectively, with longer peptides bulging out of the groove^8^. Peptide binding specificity is restricted by allotype because the amino acids of peptide anchor residues complement those in the HLA groove, leading to allele-specific sequence motifs. Meanwhile, the amino acid propensities of non-anchor positions are generally more permissive which, when combined with the HLA’s polymorphic nature, leads to a large combinatorial complexity in the number of potential pHLA structures. As a result, estimating pHLA binding affinities has been resolved using sequence data alone^9^. Predicting peptide immunogenicity and cross-reactivity, on the other hand, necessitates detailed structural information which has been more challenging to model due to subtle structural changes which may arise from amino acid variations^10^.

The high degree of structural conservation among pHLA complexes suggests that structure-based modeling of novel peptide/HLA antigens can be addressed using conventional comparative modeling approaches^11, 12^. Following the first crystallographic structure determination of a peptide/HLA complex^13^, we now have access to a substantial dataset of reliable structures^14^. Recent efforts in *de novo* modeling of the pHLA complex have focused on three main docking strategies: constrained backbone, constrained termini, and incremental peptide reconstruction^15^. Constrained backbone based predictions assume that all backbones of the same length are similar in their conformation^12, 16–21^. On the other hand, constrained termini methods offer a less restrictive view as they assume that termini residues are confined to defined “pockets” in the peptide-binding groove while the locations of the other peptide residues are said to be variable^22–27^. Lastly, fragment-based docking strategies utilize minimization protocols to sample the peptide backbone *ab initio*, allowing for a greater degree of flexibility^28, 29^. In all three cases, the modeled structures are ranked and compared based on either peptide backbone Root Mean Square Deviation (RMSD) or all-atom RMSD. While these methods have generally been able to accurately model the HLA groove and N- and C-termini of the peptide, they often miss critical details in the center of the peptide. The sequence and conformational diversity offered by the central region of the peptide defines the immunologically important area of pHLA structures^30–32^. Thus, accurate modeling of the center of the peptide is difficult but necessary to understand and predict the molecular basis for immunogenicity.

Here, we develop a database of pHLA structures (HLA3DB) and classify peptide backbones based on the sequence separation between their primary anchor residues. Focusing on nonameric peptides, we establish an internal coordinate-based system to compare backbones in dihedral angle space, and find that HLA-I peptides can sample a discrete set of conformations. Additionally, we observe that similar backbones can be obtained despite dissimilar peptide and HLA sequences, revealing the complexity of convergent interactions which ultimately define the peptide backbone. Exhaustive modeling simulations using Rosetta reveal that distinct peptide backbones necessitate unique biases of central peptide sequences, arising from several factors, including steric hindrance. Finally, we combine our basis set of distinct structural templates with a regression model trained on Rosetta energy terms to develop a structural modeling approach for nonamer/HLA complexes (RepPred). Using a cross-validation technique, we find that our method outperforms six state-of-the-art methods^24, 25, 29, 33–35^ showing a 19% accuracy improvement relative to the top method^35^, while consistently identifying the correct templates for target backbones that sparsely populated in the PDB. Additionally, independent testing using a blind set of targets shows comparable accuracy as seen in our cross-validation benchmarking results. Our findings enable accurate modeling of peptide/HLA structures at scale, paving the way for accurate prediction of peptide immunogenicity and cross-reactivity.

## Results

### Analysis of pHLA structures uncovers the basis for peptide conformational diversity

A comprehensive structural analysis of pHLA complexes requires a curated database of high-resolution X-ray structures. While there are publicly available databases that store MHC-I structural data, they do not provide a consistent format needed for further automated analysis^36–44^. Using the RCSB Protein Data Bank (PDB) Search API, we developed an automatic protocol for extracting and annotating pHLA structures with peptide lengths from 8 to 10 residues (Supplementary Fig. 1 and Methods). From each PDB entry, we retained the α_1_ and α_2_ domains of the MHC heavy chain, referred to as the MHC platform, in addition to the peptide antigen. Leveraging the IPD-IMGT/HLA Database^3^ as a reference, we then assigned an HLA allotype to each human MHC-I X-ray structure. The resulting set of curated pHLA platform structures were stored in HLA3DB (https://hla3db.research.chop.edu), a publicly available, auto-updating database. HLA3DB consists of 393 structures with 15 octamer, 296 nonamer, and 82 decamer peptides (Fig. 1A). In terms of HLA peptide binding specificities, the classical MHC-Ia allotypes present in the database cover five out of six known HLA-A and all HLA-B supertypes^45^, in addition to non-classical type Ib complexes (HLA-G, HLA-E, HLA-F). As expected, the A02 supertype is represented by 42% of the structures in the dataset. Notwithstanding, the wide range of HLA groove and peptide sequences present in our dataset provides a comprehensive sampling of the conformational space covering possible peptide backbones to guide structural modeling efforts.

**Figure 1.**
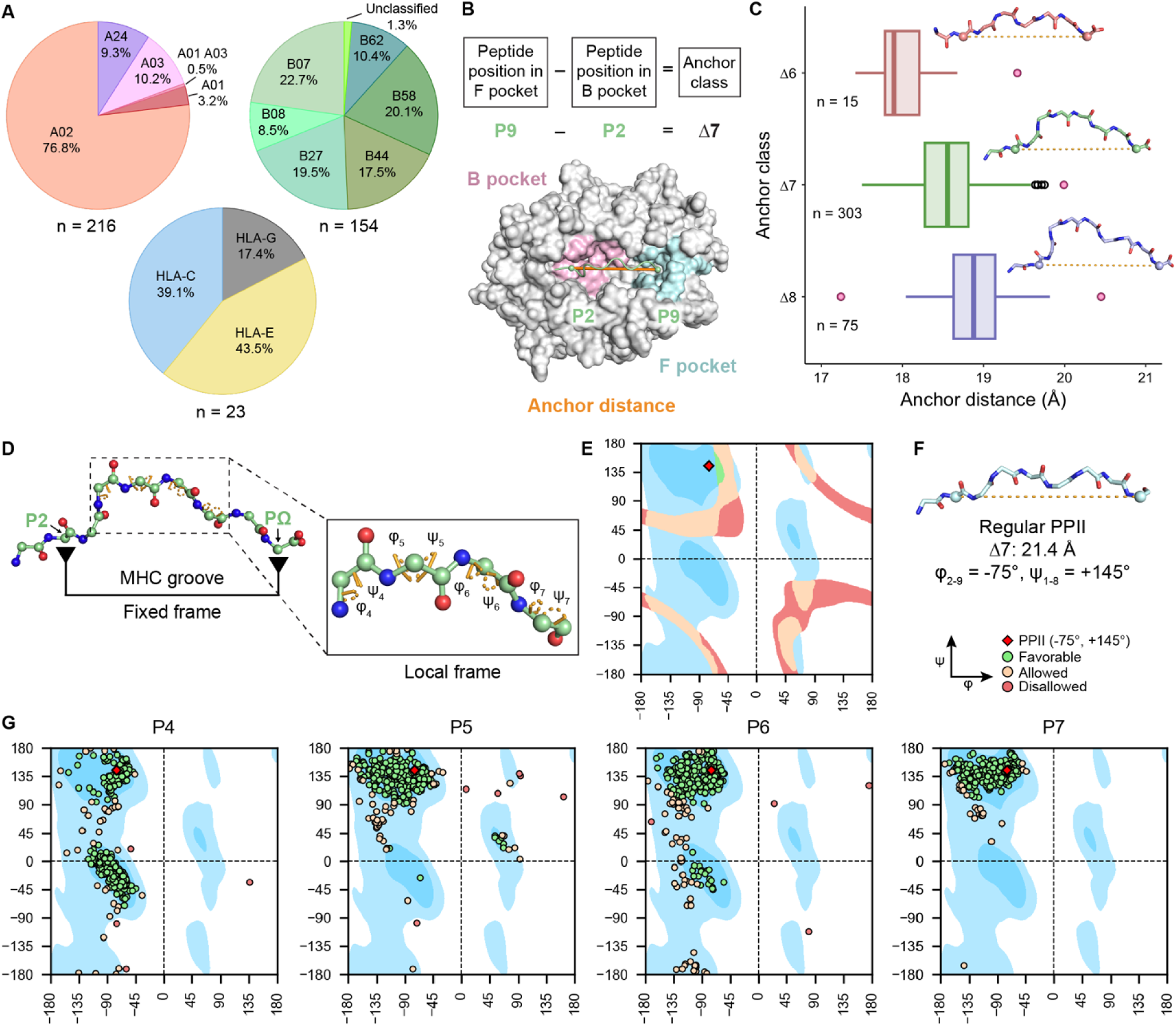
Analysis of pHLA structures in HLA3DB uncovers the basis for peptide conformational diversity. **(A)** Distribution of pHLA structures in HLA3DB by supertype for HLA-A and HLA-B and by single-resolution allotype for HLA-C, HLA-E, and HLA-G. **(B)** Schematic depicting how anchor class and anchor distance is defined using the Δ7 anchor class as an example. The Cα atoms of anchor residues are shown as green spheres and are connected by an orange solid line depicting the anchor distance. The B and F pockets are labeled in pink and blue, respectively and the remainder of the HLA is colored in grey. **(C)** Distribution of anchor distances for each anchor class. Whiskers extend to the furthest values that lie within the 75th and 25th percentile value ±1.5 times the interquartile range. Outliers are shown in black and pink circles with pink data points elaborated in Extended Data Fig. 1C–F. Peptide backbones corresponding to the median anchor distance of each anchor class are shown above each respective boxplot (Δ6: PDB ID 1E28, Δ7: 1K5N, Δ8: 3I6K). **(D)** A schematic of the fixed-local framework for conformational diversity. On the right, the inset highlights the central bulge of the peptide and the respective dihedral angles in orange dashed sectors. Backbone heavy atoms are shown as spheres. **(E)** General Ramachandran plot showing dihedral angle pairs satisfying the anchor distance distribution seen in the Δ7 anchor class (favorable overlaps are colored green, allowed overlaps are colored cream, and disallowed overlaps are colored red). The remaining regions of the Ramachandran plot are colored as favorable regions shaded blue, allowed regions shaded light blue, and unfavorable regions shaded white. The PPII conformation is shown as a red diamond at (−75°, +145°). **(F)** Backbone of a *de novo* designed polyglycine nonamer with all dihedral angles set to the PPII angle (−75°, +145°). The Cα atoms of anchor positions are shown as spheres and connected via an orange dotted line to indicate anchor distance. **(G)** General Ramachandran plots of all Δ7 nonamers (n = 293). Plots are shaded identically to panel (B) with discrete points instead of shaded surfaces.

The prominent peptide classification scheme for immunopeptidomics analysis utilizes a linear view of epitope lengths, where for most HLA allotypes the peptide’s second (P2) and last (PΩ) position bind to the B and F pockets of the peptide-binding groove, respectively. However, notable exceptions exist including HLA-A*02:01 bound to the MART-1 nonamer antigen (LAGIGILTV) which has non-canonical anchor residues at P1 and PΩ (Extended Data Fig. 1A). This exception has important ramifications from a structural perspective, indicating that peptide configurations of the same length can deviate significantly. Thus, to classify peptides of different lengths according to a global frame of reference defined by the HLA groove, we employed an anchor residue-based scheme in which an anchor class is defined as the sequence separation between the two most distant anchor residues (Fig. 1B and Methods). Using this scheme, we classified all pHLA structures in HLA3DB and found three distinct classes of peptides: Δ6, Δ7, and Δ8. Generally, octamer, nonamer, and decamer peptides reside in the Δ6, Δ7, and Δ8 classes, respectively, but this is not always the case (Extended Data Fig. 1B). As expected, the Δ7 anchor class contains the most structures, as nonameric peptides are the dominant sequences presented by MHC-I molecules. Next, we evaluate the distance between the Cα atoms of the anchor residues for each anchor class, termed the anchor distance. Despite differences in peptide-binding motifs across HLA supertypes, we find that the anchor distance is heavily confined by the geometry of the HLA groove. Examination of pHLA structures reveals overlapping distributions of anchor distances, with a median of 18 Å, 18.5 Å and 19 Å, for the Δ6, Δ7, and Δ8 classes, respectively (Fig. 1C). As the distance separation between anchor residues increases, the peptide’s central bulge becomes more pronounced, resulting in a more concave structure. A small set of outliers can be attained through either β-strand extended (Extended Data Fig. 1C, D, F) or α-helical condensed peptide backbone conformations (Extended Data Fig. 1E). In general, peptides belonging to the same anchor class show a relatively restricted distance distribution, supporting a theme where divergent conformations are attained through local changes in internal backbone degrees of freedom, rather than through global changes in the overall length of the displayed peptide antigen.

We next sought to characterize the conformational diversity of peptides belonging to the same overall anchor class. Thus, we focused on Δ7 nonamers (n = 293) as they were the most common classification in our dataset. While these peptides exhibited a narrow anchor distance distribution from 17.5 Å to 20 Å, highly divergent conformations were observed, which could influence recognition by T-cell receptors (TCRs). Building on the knowledge that TCRs preferentially interact with the central bulge of the peptide, we employed a structural framework, termed the fixed-local frame, to capture peptide conformational diversity (Fig. 1D). We establish that the primary anchor residues (P2 and P9) provide a fixed frame of reference by defining a narrow range of allowed Cα-Cα distances. Concurrently, the central portion of the peptide, specified as positions 4 to 7, define peptide backbone diversity through changes in the φ and ψ dihedral angles.

To further highlight that dihedral angle variations occurring at specific peptide positions are required to fit a nonamer in the peptide-binding groove, we modeled different regular backbone structures *de novo* in the absence of the MHC, and measured the resulting Cα-Cα distances between the primary peptide anchor residues, P2 and PΩ (Methods). Peptide anchor distances within the observed range for pHLA structures from the Δ7 anchor class (17.5 Å to 20.0 Å) were plotted as continuous surfaces on a Ramachandran plot (Fig. 1E). We found a single overlapping area of this surface with the favorable region of a Ramachandran plot that was adjacent to the ideal polyproline type II (PPII) helical conformation (φ = −75°, ψ = +145°)^46^. However, the anchor distance of an ideal PPII peptide was found to be 21.4 Å, lying outside of the distribution observed for the Δ7 anchor class (Fig. 1F). Notably, even pHLA complexes with a near regular PPII peptide conformation required slight variations to allow for an acceptable anchor distance distribution (Extended Data Fig. 2). This calculation reveals that, in order to satisfy the observed range of anchor distances in the Δ7 class of peptides, individual residues within Δ7 nonamers must adopt configurations that deviate from the ideal PPII conformation, introducing structural diversity. We next analyzed the distributions of φ and ψ dihedral angles for each peptide position among the Δ7 nonamers in our database (Fig. 1G and Extended Data Fig. 3). We observed that for the majority of structures, residues P2, P3, and P8 were generally conserved and clustered around the PPII conformation. On the other hand, the central residues of the peptide (P4 to P7) showed frequent deviations from the PPII region towards a more extended (β-strand) or more condensed (α-helix) structure. The magnitude and order of the PPII divergences, driven both by the identity of the HLA groove and peptide residues, defines the conformational diversity of epitope backbones. These results support that the conserved anchor residues (P2 and PΩ), which are primarily involved in HLA binding, display little variability from the ideal PPII conformation, while changes in the central part of the peptide (P4 to P7) promote conformational diversity necessary for specific TCR recognition.

### Unbiased classification reveals conserved peptide backbones across HLA allotypes

Towards obtaining a quantitative method for measuring the degree of structural divergence between peptide conformations according to insights from our analysis of Δ7 nonamers, we established an internal coordinate-based metric to compare the φ and ψ dihedral angles of P4 to P7, termed the D-score (Methods). This is in contrast to the widely used root-mean-square deviation (RMSD) which is defined in the Cartesian space using a global frame of reference^47^. To compare the performance of D-score vs RMSD in the specific task of comparing peptide backbones, for all pairs of Δ7 nonamers, we determine the backbone heavy atom RMSD of P4 to P7 and D-score. While low D-score values generally correspond to low RMSD values, there are significant differences between peptide backbones leading to an increased D-score that are not captured by RMSD (Extended Data Fig. 4A). Thus, our results suggest that D-score more accurately captures the difference between two peptide conformations. In contrast, the global frame used by RMSD does not accurately assess variations between individual φ and ψ dihedral angles, limiting a more precise, quantitative analysis. We define the D-score criteria as a D-score of less than 1.5 as this ensure that the RMSD is less than 1 Å between backbones. An exemplar comparison of two structures reveals how significant dihedral angle differences cannot be captured by backbone heavy atom RMSD, leading to an inaccurate evaluation of structural similarity (Extended Data Fig. 4B). We find that a large dihedral angular difference at P5 can cause a substantial backbone deviation at P6 and P7 (Extended Data Fig. 4C). Nonetheless, the resulting backbone heavy atom RMSD of P4 to P7 is less than 1 Å, suggesting a near identical peptide configuration. These results establish that the D-score, a metric in an internal coordinate system, is a more accurate measure than RMSD in determining the structural similarity of the central part of peptide backbones.

To assess the extent of structural similarity among Δ7 nonamers, for each peptide, we determined the number of neighbors i.e. non-self-peptides which are within a D-score of 1.5. Overall, each structure had a median of 28 neighbors; however the number of similar backbones ranged from 0 to over 100 (Extended Data Fig. 5A). Furthermore, we generally found that two pHLA complexes could be neighbors irrespective of their peptide or allele sequence, signifying that interactions between divergent peptide and HLA residues can yield similar peptide backbone structures. To visualize this concept, we analyzed an exemplar peptide conformation in complex with HLA-C*05:01 (Extended Data Fig. 5B). This structure had 12 neighbors, each of which exhibited similar peptide backbone dihedral angles, regardless of allotype (Extended Data Fig. 5C, D). As a result, this finding confirmed that the HLA allotype, which restricts the peptide sequence at anchor residues, plays a negligible role in defining the backbone conformation and instead structural diversity is defined by non-anchor residues, specifically P4 to P7. Recognizing that the Δ7 nonamer backbones present in our dataset can adopt recurrent structures, we sought to establish a minimal set of peptide configurations which could describe the entire conformational space. Due to its limited size, our dataset does not accurately capture the frequency of different backbones which can be adopted by immunodominant peptide antigens. Thus, it was critical that peptide conformations with zero neighbors were retained. Hence, we chose a greedy algorithm based on the fixed-local frame and constructed a binary symmetric adjacency matrix, using the D-score metric to evaluate structural similarity (Fig. 2A and Methods). In this process, the structure with the most neighbors, termed the representative, was identified as a discrete peptide backbone and it, along with its neighbors, was removed from the matrix. This procedure was continued iteratively until the matrix was empty, i.e., until all structures were accounted for. In this way, we were able to capture the full peptide conformational space without redundancy while also accounting for underrepresented backbones. As per our expectation, we found that our set of 293 Δ7 nonamer pHLA structures could be represented by a minimal set of 34 peptide backbones.

**Figure 2.**
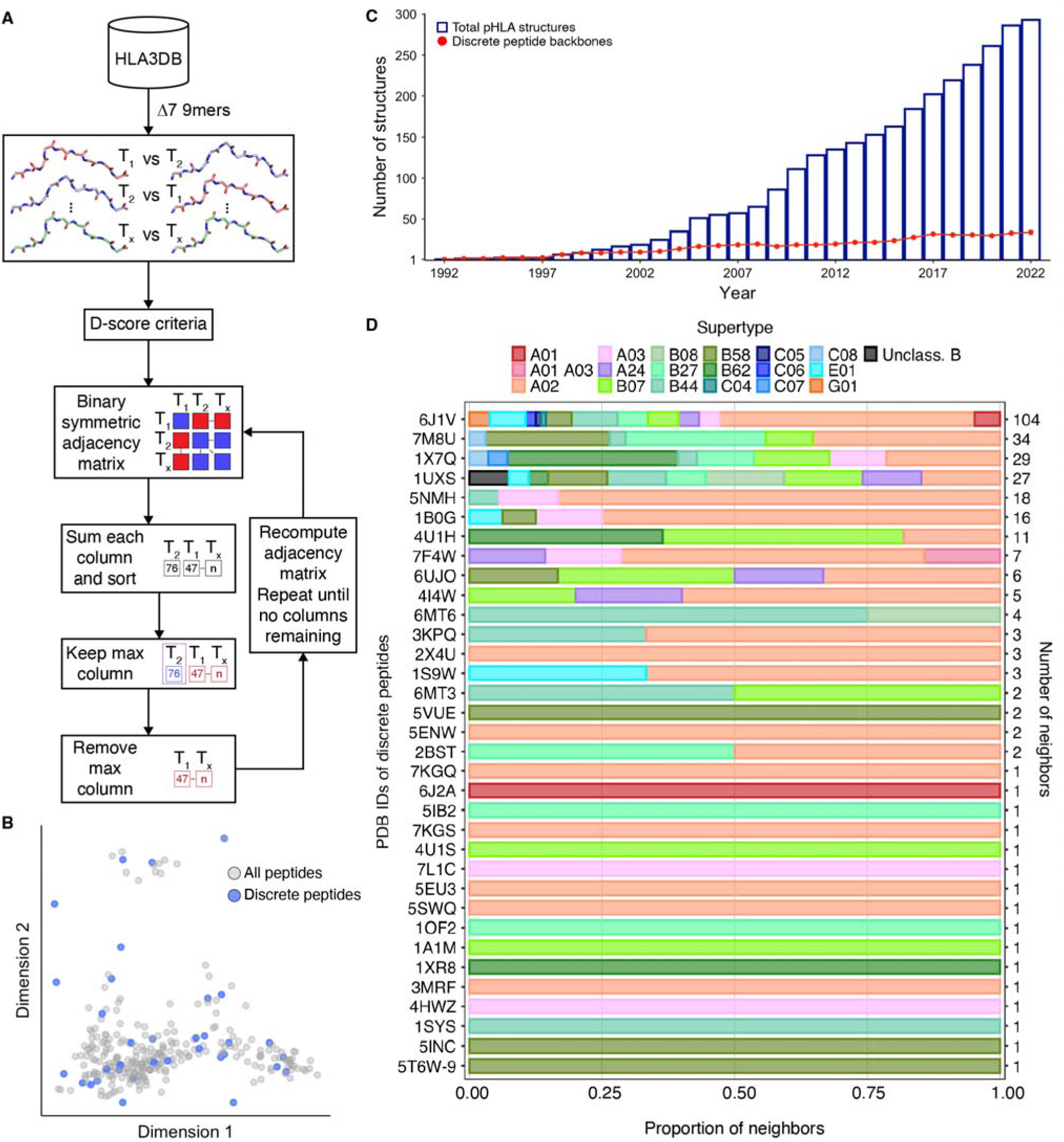
Unbiased classification reveals conserved peptide backbones across HLA allotypes. **(A)** A schematic of the greedy algorithm used to select representative peptide backbones. **(B)** A two dimensional PCA plot standardized with the sine of the dihedral angles of P4 to P7 explaining 65% of the variance. **(C)** Historical analysis of the cumulative number of structures (blue) and discrete peptide backbones (red) in HLA3DB as determined iteratively by the greedy algorithm. **(D)** The proportion of supertype in each discrete peptide’s set of neighbors. The number of neighbors for each discrete peptide is shown in the right y-axis.

To visualize the extent to which our discrete peptide backbones spanned the peptide conformational landscape, we used a two-dimensional PCA plot. This revealed that discrete peptides were not only able to capture common conformations such as those located in the lower left of the PCA plot, but also rare configurations represented by data points in the upper center region of the plot (Fig. 2B). Thus, our greedy algorithm successfully captured the entire conformational space. A historical analysis of Δ7 nonamer pHLA structures in the dataset showed a geometric progression in the number of structures deposited (Fig. 2C). Notably, the number of discrete peptide backbones has stagnated over the past five years. We next sought to understand the supertype distribution and number of neighbors for each discrete epitope. We defined a discrete peptide backbone and its neighbors as a distinct “set” of backbones. Since supertypes were generally diverse among each set of backbones, we recapitulated our finding that similar epitope conformations are not solely defined by the groove residues, but rather a convergent set of interactions between HLA and peptide residues (Fig. 2D). Furthermore, the most common peptide backbone (PDB ID 6J1V), accounted for 35% of all backbones in our dataset. In contrast, just under half of the discrete peptides defined unique classifications with just one neighbor. These results indicate that our basis set of 34 representatives can capture the diversity of pHLA structures in our dataset, and that the existing structural data likely cover the majority of possible Δ7 peptide backbone conformations.

### Exhaustive enumeration of the peptide sequence space reveals biases across discrete backbones

Next, we sought to investigate if peptide sequence biases existed across distinct backbones. However, our dataset dwarfed the potential sequence space of 9^20^ peptides, containing just 247 nonredundant Δ7 nonamer sequences. Thus, we expanded our existing peptide sequence space coverage using Rosetta modeling simulations (Fig. 3A and Methods). To ensure that the peptide-binding groove was not a confounding variable, we restricted our analysis to HLA-A*02:01, the most common allotype in our dataset. Using the aforementioned greedy algorithm, we determined that there were 20 distinct peptide backbones across all 121 HLA-A*02:01 Δ7 nonamers. We focused our analysis on the five most common HLA-A*02:01 configurations, determined by their number of neighboring conformations. Next, we narrowed down the set of potential peptide sequences by applying two constraints. First, along with the center of the peptide (P4 to P7), P3 and P8 were also included to capture any effects that adjacent positions may have on the peptide sequence of the central residues. Second, we leveraged known sequence biases from eluted peptides in the Immune Epitope Database (IEDB)^48^ for P3 to P8 to narrow our sequence space from 20 amino acids to 8 to 11 amino acids, depending on the position (Supplementary Table 1). We also considered proline at each position due to its restraining effects on the peptide backbone. In total, we evaluated 784,080 peptide sequences for each of the five distinct peptide backbones. Iterating through the set of peptide sequences, we generated models, determined the total score of each model using the ref2015 Rosetta energy function^49^, and only considered the peptide sequences of models that were in the top one percent of the energy distribution, i.e., 7,840 models. As a result, we substantially expanded our dataset of HLA-A*02:01 epitopes while accounting for the constraints placed on the peptide sequence space by the groove.

**Figure 3.**
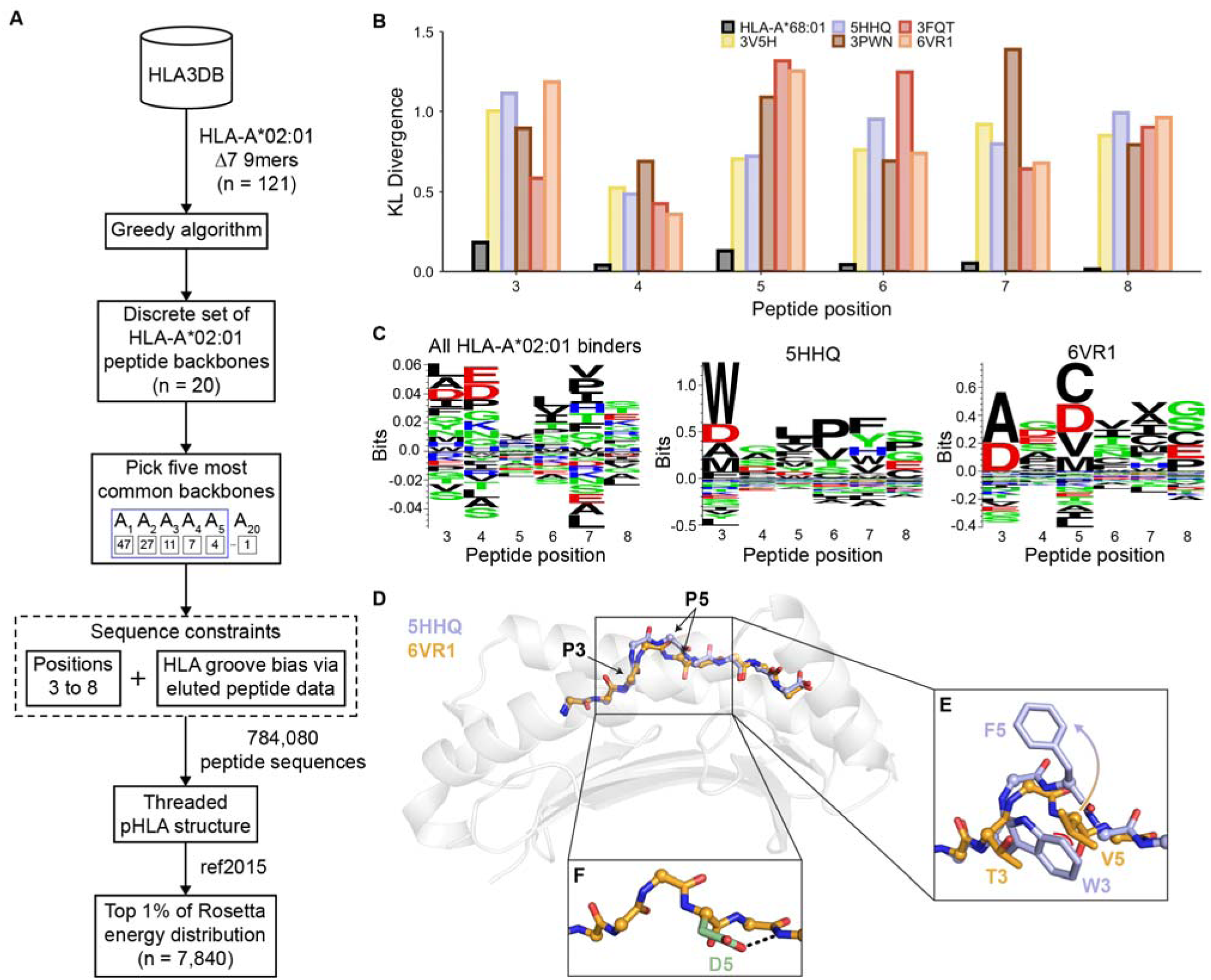
Exhaustive enumeration of peptide sequence space reveals biases across backbones. **(A)** A schematic of the exhaustive structural modeling conducted to augment the existing peptide sequence space for HLA-A*02:01. **(B)** Kullback-Leibler (KL) divergence between each of the predicted sequence space of the five most common HLA-A*02:01 backbones represented by PDB IDs and HLA-A*02:01 reference sequence space. A comparison between the eluted peptides bound to HLA-A*68:01 and those bound to HLA-A*02:01 was included as a negative control. **(C)** Peptide sequence logos of all eluted peptides shown to bind to HLA-A*02:01 via the IEDB (n = 32,483) and from structural modeling results (n = 7,840). Created using Seq2Logo^80^. **(D)** Structural overlay of the peptide backbones (5HHQ, blue and 6VR1, orange) bound to HLA-A*02:01 (gray) with the peptide Cα atoms shown as spheres. **(E)** An enlarged view of the middle of the peptide with side chains depicted for P3 and P5. Coloring of the peptide backbone and side chain is identical to panel (D). Steric hindrance is shown as a red curve. **(F)** Structural model of the 6VR1 peptide backbone with P5 mutated to aspartate. A hydrogen bond is shown as a dashed black line.

To determine if these backbone-specific peptide sequences were distinct from the baseline peptide binding preferences, we computed the Kullback–Leibler (KL) divergence^48^ at each position using 32,483 peptides experimentally verified to bind to HLA-A*02:01. We utilized peptide binding data from HLA-A*68:01, an allele belonging to the A02 supertype and therefore showing a similar peptide binding specificity, as a measure of the lower bound of the KL divergence. Our results revealed that all five distinct backbones caused peptide sequence biases (Fig. 3B). While the KL divergence was consistently lower for P4 compared to other positions, the HLA-A*02:01 binding motif contained a known bias^50^ for negatively charged amino acids at P4, ultimately decreasing the perceived sequence bias. We confirmed the quantitative values given by the KL divergence by visually comparing the sequence logo of all known HLA-A*02:01 binders to those obtained by exhaustive structural modeling (Fig. 3C and Extended Data Fig. 6A, B, and C). We next sought to explain why different backbones could lead to distinct peptide sequence biases. As an example, we chose two representative peptide configurations (corresponding to PDB IDs 5HHQ and 6VR1) with a D-score of 5.9 (Fig. 3D). A per-position D-score analysis revealed that the primary deviation occurred at the ψ_6_ angle (Extended Data Fig. 6D). However, the backbones appeared to diverge prior to P6, suggesting that compensatory backbone motions were required to accommodate for this dihedral angle difference. Peptide sequence logos of these backbones also revealed significant differences. For 5HHQ’s backbone, our modeling approach reported a preference for tryptophan at P3 which was also found in the native peptide sequence (GIWGFVFTL). Meanwhile, the sequence logo of 6VR1 favored small amino acids at P3 such as alanine since a bulky group at this position would clash with most side chains at P5 (Fig. 3E). The native peptide sequence of 6VR1, HMTEVVRRC, followed this trend as well. A backbone deviation in 5HHQ diverted the side chain at P5 away from the tryptophan, avoiding an energetically unfavorable steric hindrance issue and encouraging a broad selection of sequences. Additionally, we found a preference for aspartate at P5 in the sequence logo of 6VR1 which can be explained by a favorable hydrogen bonding interaction occurring between the side chain and backbone (Fig. 3F). Due to 5HHQ’s backbone conformation, the side chain of P5 cannot contact the backbone nitrogen atom and thus does not display the same sequence bias. This comparison demonstrates how the backbone conformation could impose local steric constraints or introduce favorable interactions through crosstalk between peptide backbone and side chain features. Also, it suggests that long-range interactions could similarly occur between the backbone and HLA groove residues and cause dissimilar pHLA combinations to adopt the same configuration.

### A trained regression function allows for accurate structural modeling

We aimed to harness the knowledge that all pHLA structures in our dataset could be captured by a set of representative backbones to enable accurate structural modeling of new peptides on all HLA allotypes. To this end, we developed RepPred, an automated modeling method which utilizes discrete backbones as templates for homology modeling of Δ7 nonamer epitopes (Fig. 4A). Briefly, a target peptide sequence is threaded onto 33 stable representative templates, followed by structural refinement in Rosetta to create an initial set of models (Methods). We computed the per residue Rosetta energy terms of the models^29^ and utilized these values as input features for a support vector machine regression (SVR) function, enabling the prediction of the D-score between the native crystal structure and the model. RepPred reports the best model as that with the lowest predicted D-score.

**Figure 4.**
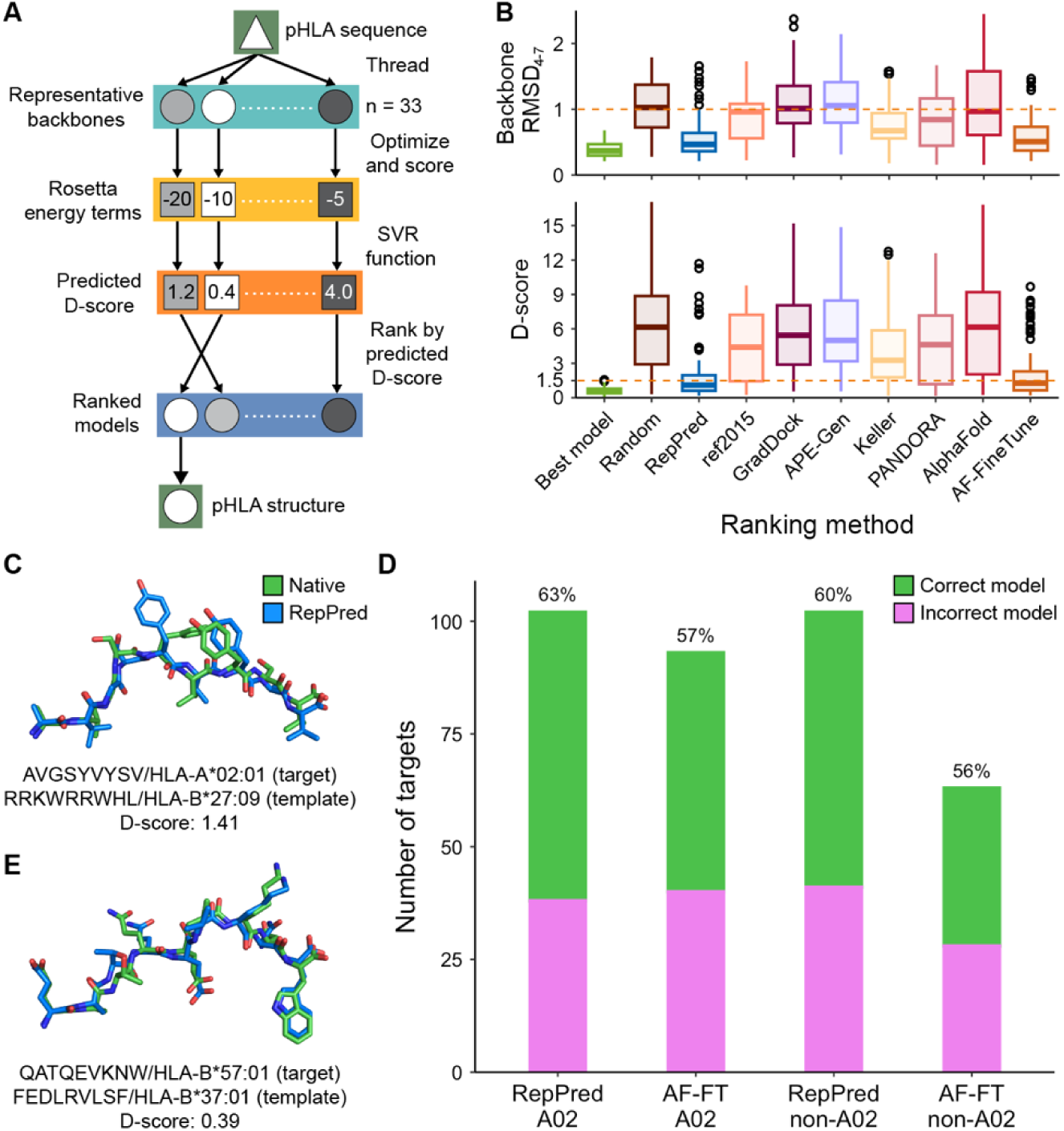
Structural modeling of nonamer/HLA complexes (RepPred) and comparison to state-of-the-art methods. **(A)** Full workflow of the RepPred structural modeling method. **(B)** Boxplots showing the distribution of the D-score and backbone heavy atom RMSD of RepPred and state-of-the-art methods, sorted by publication date. Whiskers extend to the furthest values that lie within the 75th and 25th percentile value ±1.5 times the interquartile range and outliers are shown in black circles. An orange dashed line is at a D-score of 1.5 and RMSD of 1.0. **(C)** Structural superposition of the RepPred model (template PDB ID 1OF2) and native structure (PDB ID 7U21) of the blind target melanoma antigen AVGSYVYSV bound to HLA-A*02:01. The target sequence and allotype is written below the structure followed by the sequence and allotype of the template. **(D)** Comparison of RepPred against AF-FT for A02 and non-A02 targets. Accuracy is shown as a percentage over each bar. **(E)** Structural superposition of the RepPred model (template PDB ID 6MT6) and native structure (PDB ID 7R7Y) of the blind target HIV antigen QATQEVKNW bound to HLA-A*57:01. Formatting is identical to panel (C).

To assess the efficacy of RepPred, we perform a benchmark on all targets in our dataset against their nonhomologous representatives (see the Methods). Thus, we evaluate a total of 7,775 target-template pairs in a leave-one-out cross-validated manner whereby the models of the target structure are removed from the training set. Recognizing the overrepresentation of HLA-A*02:01 structures in our dataset, we first explore the accuracy for HLA-A02 targets separately from all other allotypes i.e., non-A02, targets. RepPred reports a structurally accurate model for 63% of A02 targets (n = 102) and, on the basis of backbone RMSD for the middle of the peptide, it selects a sub-angstrom model for 92% of A02 targets (Fig. 4B). While the refinement step can cause the backbone to drift from the template conformation, selecting the best model for A02 targets by D-score gives an accuracy of 99% which is generally in agreement with the finding that the representatives cover the conformational space. On the other hand, a random selection of the representative backbone rarely (10%) selects the correct decoy, as expected. To validate our modeling approach, we conduct a blind test by assessing the accuracy of RepPred on five Δ7 nonamer/HLA-A*02:01 structures in HLA3DB deposited after the cutoff date for our dataset. In line with the benchmark results, we accurately predict three out of five structures within a D-score of 1.5, allowing for accurate placement of side chains in their native rotameric states (Extended Data Fig. 7A). Our blind test included a melanoma neoantigen (AVGSYVYSV)^51^ which we modeled with a template structure originating from HLA-B*27:09, an allele not found in the target’s supertype family (Fig. 4C). This finding recapitulates the trend that similar backbones can occur independent of allele identity, allowing us to use all observed backbones as a basis set for structural modeling.

Next, we compare RepPred with six existing approaches for pHLA-I structural modeling: GradDock^24^, APE-Gen^33^, a method by Keller, et al.^29^, PANDORA^25^, AlphaFold2^34^, and a peptide/MHC fine-tuned version of AlphaFold (AF-FT)^35^. We focus our comparison on HLA-A02 targets and find that RepPred outperforms five of the six methods by at least 32% with respect to D-score (Fig. 4B). We recognize that nearly half of the A02 targets are neighbors to a distinct representative backbone. Hence, a naive method could achieve 50% accuracy by simply adopting this backbone conformation for all targets. While RepPred does achieve higher accuracy (76%) for the most common backbone classification (6J1V), it maintains 50% accuracy for all other backbones, resulting in an overall accuracy of 63% (Extended Data Fig. 7B). Meanwhile, AF-FT reports an overall accuracy of 57% and an accuracy of 78% and 38% for the most common and all other backbones, respectively. Thus, while AF-FT is influenced by the bias of the most common backbone, in utilizing representative backbones, RepPred weighs each backbone equally and reduces the effect of this bias. As a result, RepPred shows a slight improvement across all backbones and a notable enhancement for less common conformations. We next sought to conduct a similar comparison of RepPred to AF-FT for non-A02 targets. RepPred achieves 60% accuracy for these targets (Fig. 4D), an accuracy comparable to that of A02 targets. We validated our method through a blind test of seven non-A02 targets in which we obtained an accuracy of 57% (Extended Data Fig. 7A). RepPred models an immunodominant HIV epitope (QATQEVKNW)^52^ with near-native side chain placement as a result of high-fidelity generation of the peptide backbone (Fig. 4E). A comparison to AF-FT reveals an overall accuracy of 56% for non-A02 targets. When assessing performance for both methods based on backbone conformation, we observe that RepPred performs 19% better than AF-FT for common backbones and comparably for all other conformations (Extended Data Fig. 7C). Taken together, RepPred models nonamer pHLA complexes with high accuracy, a finding which resurfaces in blind tests of immunologically relevant antigens, and performs better than state-of-the-art methods.

## Discussion

Our results characterize peptide backbone diversity across all pHLA-I structures. Using HLA3DB, our database of peptide/HLA structures, we broadly categorize peptides using our anchor classification scheme which accounts for noncanonical anchor residues. When combined with a comprehensive analysis of the backbone dihedral angles, we describe a framework to explain conformational diversity and introduce D-score as a measure of structural similarity. Using this metric, we find that peptide backbone similarity is allotype-independent and identify 34 discrete peptide configurations which cover the entire seen conformational landscape. Focusing on HLA-A*02:01, we discover strong peptide sequence trends influenced by distinct backbone features. Finally, we introduce RepPred, an accurate, pan-allelic structural modeling approach for nonamer/HLA complexes, and demonstrate its improved accuracy over existing methods.

There are several important limitations to our study. First, our dataset of X-ray crystal structures is not necessarily a representative sample of *in vivo* nonamer pHLA structures and thus the true distribution of backbone conformations is currently unknown. Additionally, due to the inherent bias in the motivations behind the determination of each crystal structure, our dataset likely contains a high proportion of disease-relevant peptides. Nonetheless, we are confident that we have captured a large fraction of known backbone conformations, as evidenced by our historical analysis of the PDB (Fig. 2C). Additionally, as peptide/HLA structures are solved, our automated modeling method will progressively improve due to greater sampling of underrepresented backbone conformations. Second, in utilizing crystallographic structures, we do not account for the dynamic nature of the pHLA complex which has been shown to impact TCR recognition in some cases^53^. For instance, peptide backbones have been observed to exhibit rigid body motions upon TCR binding and thus could acquire conformations not covered by our discrete backbones^54–56^.

HLA3DB contains discrete peptide backbone conformations which were utilized for structural modeling of peptide/HLA-I complexes via RepPred. Going forward, this approach can be improved by training the regression function on data from non-human or class II peptide/MHC structures and incorporating multiple sequence alignment information. Given that the peptide backbone can be accurately modeled, this structural information can be incorporated into existing TCR:pMHC modeling approaches to allow for improved prediction of binding specificity^57–60^. Additionally, RepPred can be utilized to aid the design of chimeric HLA molecules and subsequently identify peptide-centric receptors^61, 62^. Finally, leveraging structures in HLA3DB, we can identify cross-reactive peptide sequences which, when combined with large-scale structure prediction, can help to finetune immune receptors^10, 63–68^. Collectively, HLA3DB provides an atlas of peptide backbones which RepPred utilizes to traverse from the sequence to the structure space, offering an approach for predicting peptide cross-reactivity in a backbone-focused manner.

## Methods

### HLA3DB curation

The peptide/HLA-I structural dataset (HLA3DB) was curated using a custom Python script (Supplementary Fig. 1). Complexes were identified using the RCSB PDB Search API^69^ via a JSON file with the following broad criteria: (i) macromolecular name containing keywords “MHC” or “HLA” or gene name is “HLA-A”, “HLA-B”, “HLA-C”, “HLA-E”, or “HLA-G” (ii) source organism is “Homo sapiens”, (iii) structure resolution is 3.0 Å or higher, and (iv) structure release date is January 1^st^, 1988, or later. Using these selection criteria, we obtain a total of 1017 PDB entries (as of April 29^th^, 2022). Each structure is automatically fetched, filtered and saved into the database using the Bio.PDB Python package from Biopython^70, 71^. Next, we check the PDB file for readability, classify chains into HLA heavy chain, β−2 microglobulin, peptide and other depending on the sequence alignment (with >50% identity) to a reference HLA-A*02:01 heavy chain (consisting of 180 residues from N-terminus), β−2 microglobulin and sequence length (for peptides of length 8 to 10 amino acids). If the PDB file has other chains not within 5 Å from the peptide, the entry is retained for subsequent filtering. Complexes with missing N-terminal residues in the heavy chain are subjected to RosettaRemodel^72^ where missing one (glycine) or two residues (glycine and serine) from the reference HLA-A*02:01 are modeled. Structures with HLA sequence length of less than 180 residues along the C-termini of the heavy chain are discarded. After trimming the heavy chain up to 180 residues, the structure only contains the peptide/HLA complex. Any structure missing backbone heavy atoms or contains atoms with zero occupancy along the (i) peptide are removed from the database and (ii) heavy chain are reported for manual examination. The PDB entries are then HLA typed by performing pairwise sequence alignment with ∼3000 HLA sequences obtained from the IPD-IMGT/HLA sequence database^3^. The peptide and the heavy chains are renamed, the residues are renumbered, and coordinates are saved in our final structure database. A FASTA file is generated summarizing the dataset with PDB ID, chain name, allele, structure release date, and resolution followed by the sequence of either the HLA or peptide depending on the chain name specified in the previous line. Finally, the dataset was queried for decameric peptides in the Δ7 anchor class and, if appropriate, manual truncation was conducted to create additional nonameric Δ7 peptides which were named by their original PDB ID followed by “−9”.

### Anchor-based classification

The two most distant anchor residues were established by finding the peptide residue with lowest Cα-Cα distance between residue 24 and 123 in the HLA corresponding to the B and F pockets, respectively^7^. Anchor class was determined by subtracting the two most distant anchor residues. The anchor distance was computed by assessing the Cα-Cα distance between the two most distant anchor residues which define the anchor class. All analysis was completed using custom Python scripts and PyRosetta 4.0^73^.

### Dihedral angle analysis

A free polyglycine nonamer, chosen to eliminate the impact of steric hindrance between HLA residues and peptide side chains, was created using PyMOL^74^ Builder defaulted to an α-helical conformation. Dihedral angles were set using the Python PyMOL package and iterated through −180° to +180° at 1° intervals for both φ and ψ dihedral angles. For each instance, the anchor distance was computed using the aforementioned method. Dihedral angle pairs that allowed for an anchor distance between 17.5 Å and 20.0 Å were plotted on a Ramachandran plot. Visualization of Ramachandran plots was conducted using a modification of the ramachandran 0.0.2 Python package.

### D-score structural accuracy metric

The difference between backbone dihedral angles of the same type and residue position was computed using the following equation^75^.

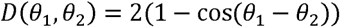

This equation accounts for the cyclic nature of dihedral angles, allowing for an accurate difference measurement. We extended this equation to determine the similarity between two Δ7 nonameric peptide backbones A and B as

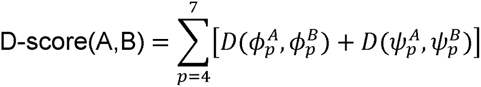

The D-score criteria states that if A and B have a D-score of ≤ 1.5, then the two backbones are considered structurally similar or neighbors.

### Classification of peptide backbones using a greedy algorithm

Backbone classification was conducted using a greedy algorithm in Python. Peptide dihedral angles of all Δ7 nonamers were computed and a binary matrix was created using the D-score criteria such that if two backbones were similar a one was added and otherwise a zero. Then, for each backbone, the number of similar structures were summed. The backbone with the most number of neighbors was noted, and it and its neighbors were removed from the matrix. This process continued iteratively until the matrix was empty.

### Exhaustive structural modeling

Before modeling peptides using pHLA structure templates, we prepared the crystal structures using the FastRelax protocol in Rosetta (Version 2020.08)^76, 77^. Then, using a Python script, we iterated through all possible combinations of amino acids between positions 3 to 8 as defined by the HLA-A*02:01 groove bias (Supplementary Table 1) and the new peptide sequence was stored in a blueprint file. The anchor residues of P2 and P9 were not included as their sequence was generally conserved across HLA-A*02:01 epitopes. Based on its distance from the center of the peptide, P1 was not included. To model different peptides onto the template pHLA structures, we utilized the RosettaRemodel^72^ application from the Rosetta suite of programs. All side chains of the template structure aside from those modeled on the peptide were left in their original poses. The Remodel application automatically scores the modeled structure and the total score of the complex was utilized. The top 1% of all structures by total Rosetta score were used to establish the peptide sequence space of a given discrete peptide backbone. Next, a position probability matrix (PPM) was created from sequence data with zeroes defaulted to 0.02. The Kullback-Leibler (KL) divergence was computed using the *rel_entr* function in SciPy^78^ with eluted HLA-A*02:01 and HLA-A*68:01 peptides from the IEDB as a reference.

### Structural modeling and regression

Prior to structural modeling, we perform a stability check on the crystal structures of the discrete peptide backbones identified by the greedy algorithm using the Cartesian relax protocol in the Rosetta forcefield. The backbones which moved more than a D-score of 1.0 upon relaxation were removed from the template set and the greedy algorithm was reapplied to ensure complete coverage of the conformational space.

We set the amino acid sequence of the nine peptide residues and 180 residues of the MHC using the *PartialThreadingMover* in PyRosetta^73^. The threaded models were then optimized using the Cartesian relax protocol and if the D-score between the template crystal structure and relaxed model was greater than 1.5, these models were completely removed. For each model produced in threading, we computed 129 per residue energy terms of peptide from the ref2015 energy function^29^ and these energy terms are further used for regression analysis. We trained a regression function to predict the D-score between the predicted models and the corresponding crystal structures of pMHCs using per residue energy terms of peptides as features. We used the *scikit*-*learn*^79^ implementation of Support vector machine regression (SVR) with radial basis function (*sklearn.svm.SVR*). The free parameters of the models, namely the distance parameter epsilon and regularization parameter C, were determined through a grid search hyperparameter scanning. The grid search covered values for epsilon and C in the range 10^-4^ to 10^4^ and the best parameter combination was determined using the coefficient of determination (R^2^) score. The input features of the SVR function were transformed using a uniform Quantile Transformer to avoid non-standard distributions. RepPred can model a single target sequence utilizing 136 Xeon 2.10 GHz cores in 30 minutes.

To benchmark RepPred, we threaded the sequence of 260 peptide/HLA-A*02:01 complexes onto 33 discrete backbone templates, removing structural models generated using a template structure that had a peptide which is different from the target peptide at three or fewer residues (homologs). Additionally, in the training set only, we removed any target-template pairs with a D-score of greater than 7. This resulted in 7,775 target-template pairs. The SVR function was subjected to a leave-one-out cross validation in which we removed all data corresponding to one target structure from the training dataset and used it as our test case. For each target the model with lowest predicted D-score from the SVR is chosen as the best structural model. To assess accuracy, the D-score between best structural model and the crystal structure of the target was calculated. Additionally, the backbone heavy atom RMSD of the middle of the peptide (P4 to P7) was computed for A02 targets. For non-A02 targets, RepPred reports a model only if the predicted D-score is less than 2.0.

### Comparison to state-of-the-art methods

In comparing RepPred against six open-source methods for pHLA structural modeling, we maintained all default parameters and followed the appropriate documentation with no change to existing code. For all peptide/HLA-A*02:01 targets, we removed structural homologs for each method except PANDORA. For AF-FT, we removed any targets which were utilized to fine-tune AlphaFold. Some structures could not be processed by existing software. Thus, we report results from 96 structures for GradDock, 108 for APE-Gen, 110 for Keller, 109 for PANDORA, 110 for AlphaFold, and 162 for AF-FT. For GradDock, 3MRE was used as the template structure as it is the highest resolution HLA-A*02:01 structure in our dataset.

### Code availability

HLA3DB can be accessed via https://hla3db.research.chop.edu. Code used for structural modeling is available on GitHub via https://github.com/titaniumsg/RepPred.

## Acknowledgments

The authors thank Dr. Andrew C. McShan for helpful discussions and Hailey Wallace from the Rosetta Commons REU. This research was supported through grants by NIAID (5R01AI143997), NIGMS (5R35GM125034), and NIDDK (5U01DK112217) to N.G.S.

## Author information

### Authors and Affiliations

**Center for Computational and Genomic Medicine, Department of Pathology and Laboratory Medicine, The Children’s Hospital of Philadelphia, Philadelphia, PA, USA**

Sagar Gupta, Santrupti Nerli, Sreeja Kutti Kandy, Glenn L. Mersky, & Nikolaos G. Sgourakis

**Department of Biochemistry and Biophysics, Perelman School of Medicine, University of Pennsylvania, Philadelphia, PA, USA**

Nikolaos G. Sgourakis

**College of Arts and Sciences, University of Pennsylvania, Philadelphia, PA, USA**

Sagar Gupta

### Contributions

S.G., S.N., and G.L.M created HLA3DB. S.G. and S.N. performed computational analysis of peptide/HLA crystal structures. S.G. developed the greedy algorithm to identify discrete peptide backbones and conducted exhaustive structural modeling. S.G. and S.K.K. developed and benchmarked RepPred. S.G. and N.G.S. wrote the paper, with feedback from all authors. N.G.S. acquired funding and supervised the project.

### Corresponding author

Correspondence to Nikolaos G. Sgourakis

### Ethics declarations

All authors declare that the research was conducted in the absence of any commercial or financial relationships that could be construed as a potential conflict of interest.

## Extended Data Figures

**Extended Data Figure 1.**
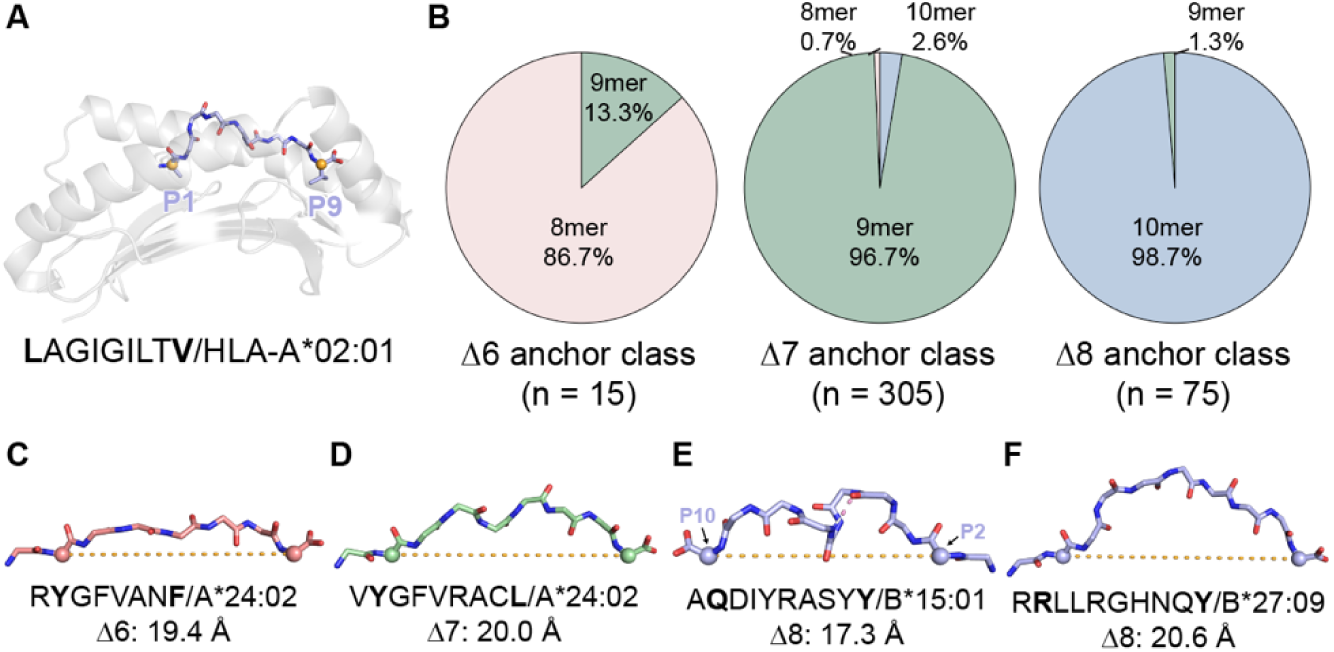
Anchor class justification and special cases. **(A)** Non-canonical binding mode of the nonameric LAGIGILTV peptide bound to HLA-A*02:01 (PDB ID 2GTW). The MHC is colored in grey and shown as cartoon while the peptide backbone is colored in blue and shown as sticks. The Cα atoms of anchor positions (labeled) are shown as orange spheres. Anchor residues are further highlighted by bold text in the peptide sequence. **(B)** Pie charts showing the distribution of peptide lengths across the three anchor classes. **(C)** An extended conformation of a Δ6 octamer peptide (PDB ID 4F7T). **(D)** An extended conformation of a Δ7 nonamer peptide (PDB ID 2BCK). **(E)** A condensed conformation of a Δ8 decamer via an interpeptide 3_10_-helix (PDB ID 5VZ5). Note that the peptide has been rotated 180°. **(F)** An extended conformation of a Δ8 decamer peptide (PDB ID 1JGD).

**Extended Data Figure 2.**
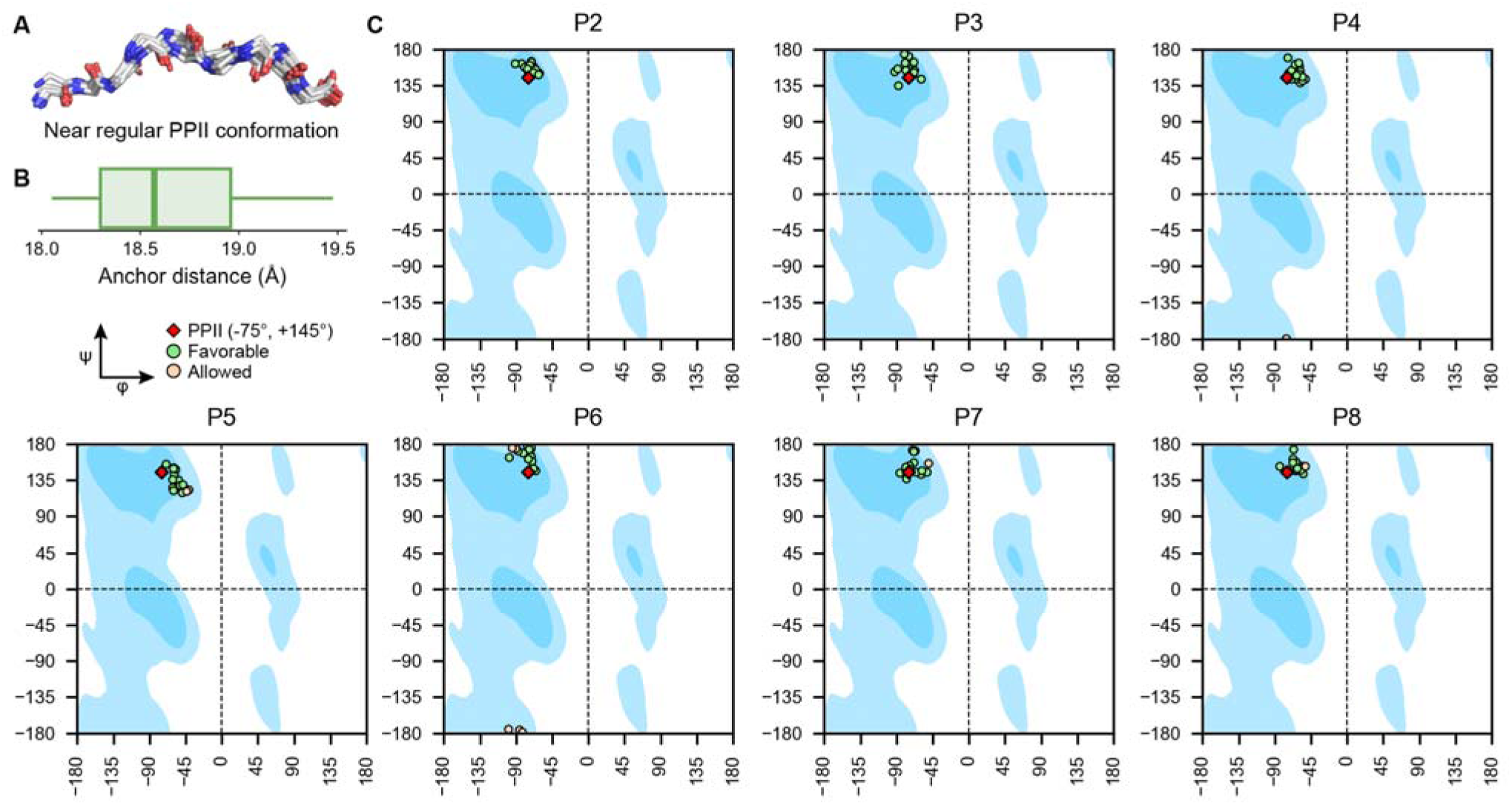
Analysis of near regular PPII backbone conformations. **(A)** Structural overlay of near regular PPII peptide backbones (n = 17). The MHC is not shown. **(B)** Anchor distance distribution of near regular PPII peptides **(C)** General Ramachandran plot showing dihedral angle pairs of the structures shown in (A). A legend is provided to the left of the plots.

**Extended Data Figure 3.**
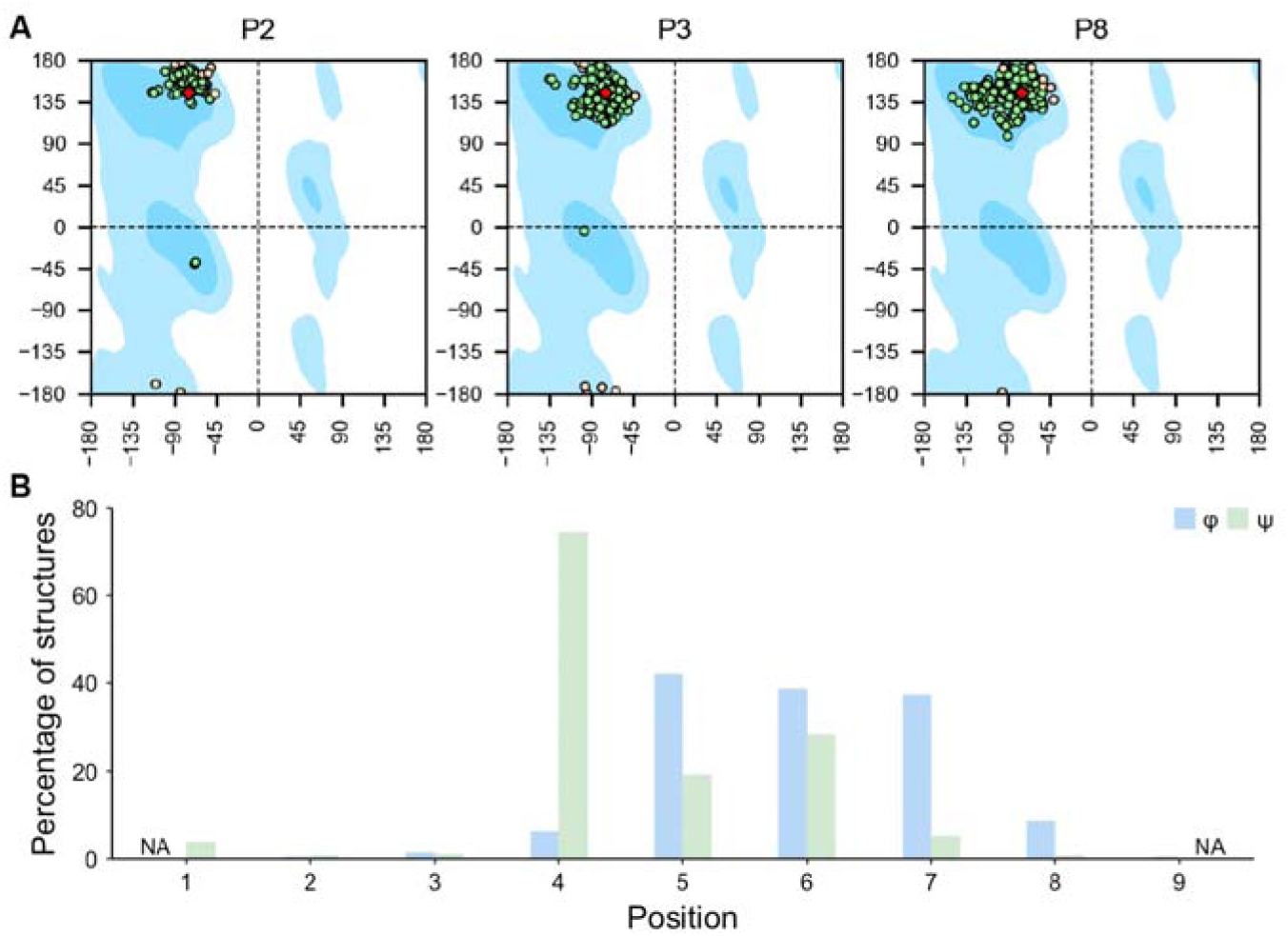
Analysis of Δ7 nonamers in HLA3DB. **(A**) General Ramachandran plots of Δ7 nonamers shaded identically to Fig. 1G. **(B)** Percentage of structures with a PPII deviation at a given dihedral angle and position.

**Extended Data Figure 4.**
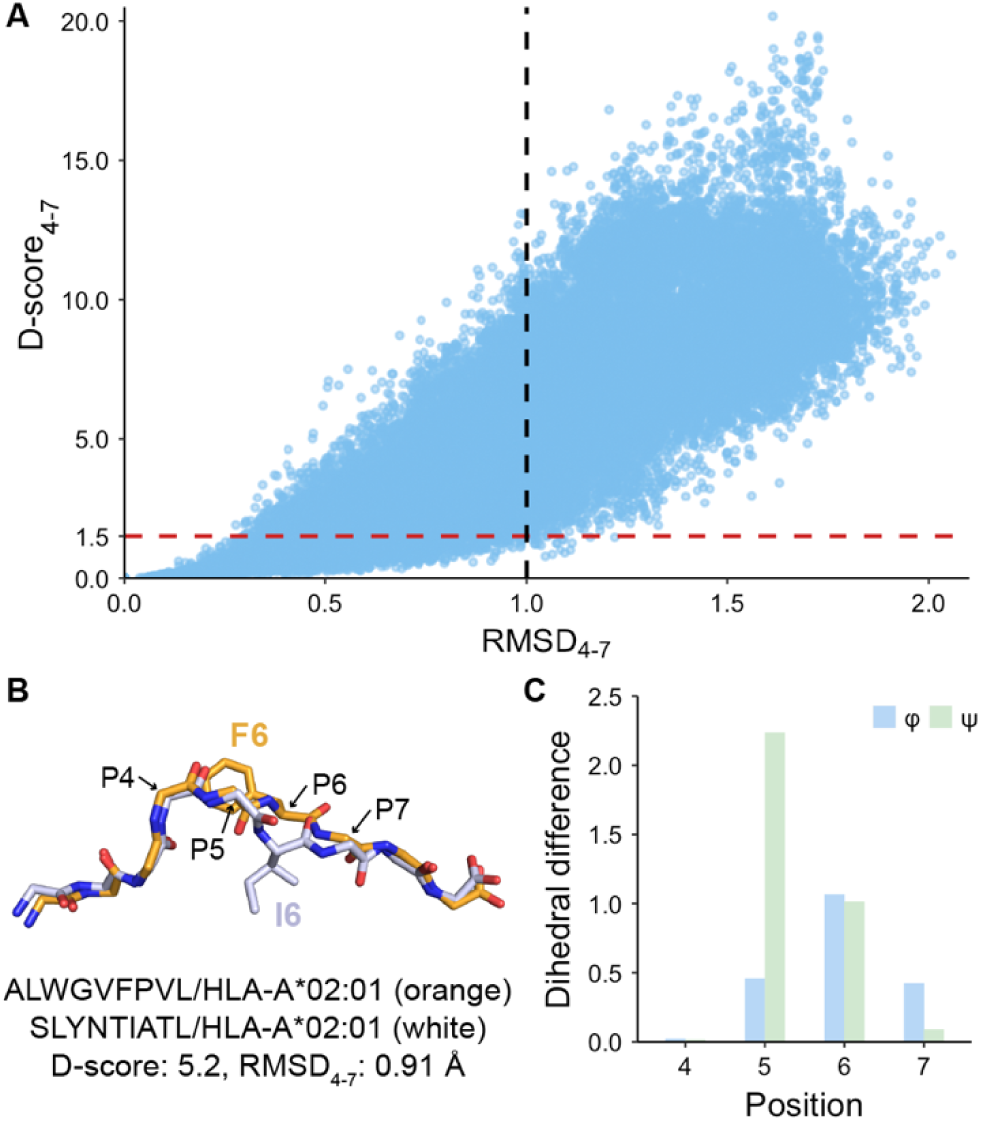
Comparison of D-score and RMSD. **(A)** Pairwise comparison of D-score and backbone heavy atom RMSD for P4 to P7. **(B)** Exemplar superposition of ALWGVFPVL/HLA-A*02:01 (PDB ID 1I7T, orange) and SLYNTIATL/HLA-A*02:01 (PDB ID 5NMH, white). The MHC is not shown. **(C)** Dihedral difference by position and angle for the exemplar structural superposition shown in (B).

**Extended Data Figure 5.**
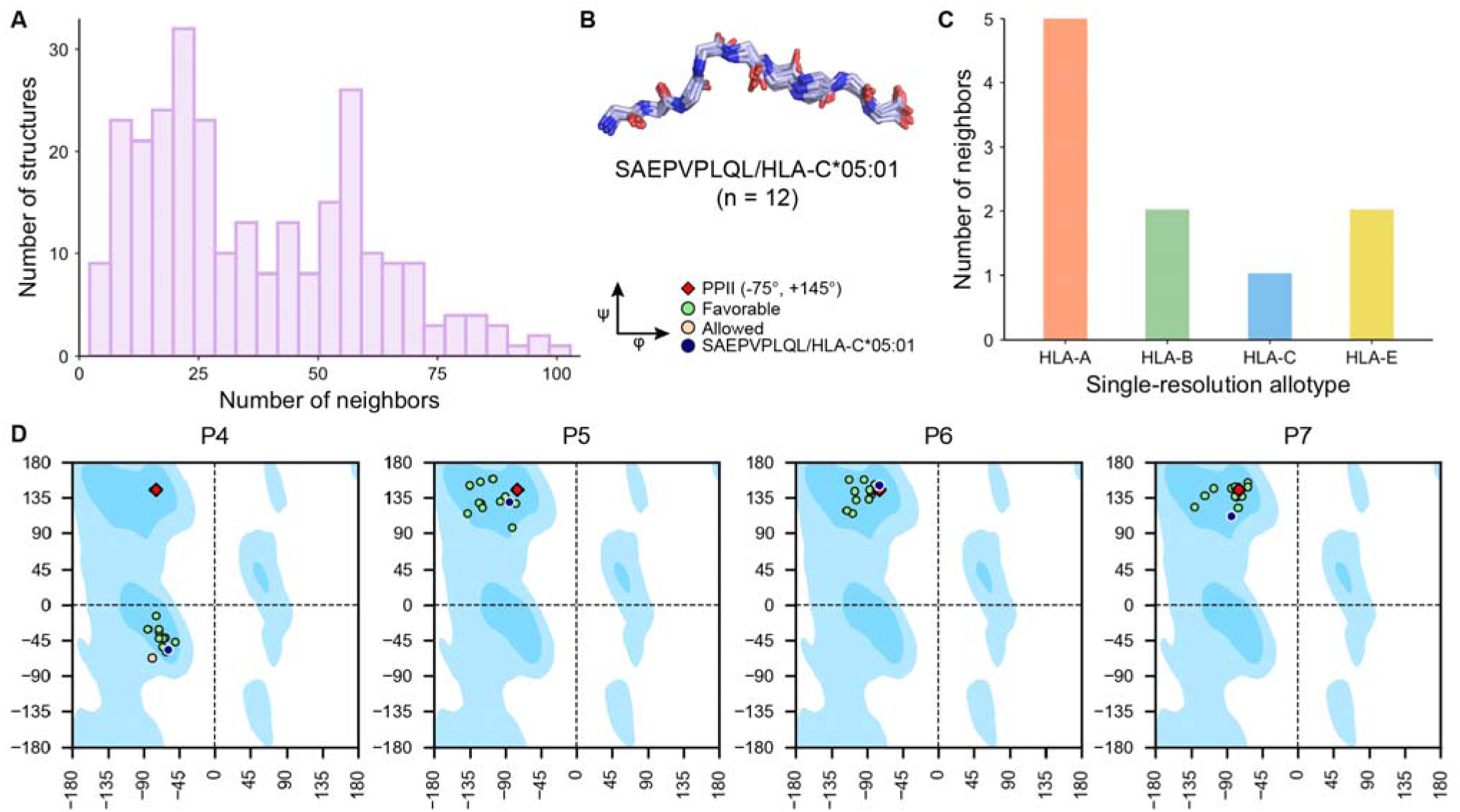
Structural conservation of peptide backbones occurs across allotypes. **(A)** Number of neighbors per Δ7 nonamers backbone. **(B)** Structural superposition of neighboring backbones to the exemplar peptide via peptide backbone heavy atoms. The MHC is not shown. **(C)** Number of neighbors of the exemplar structure by single-resolution allotype. **(D)** Ramachandran plots of P4 to P7 of the neighbors (green/cream) and exemplar structure (dark blue). All other coloring is identical to Fig. 1G.

**Extended Data Figure 6.**
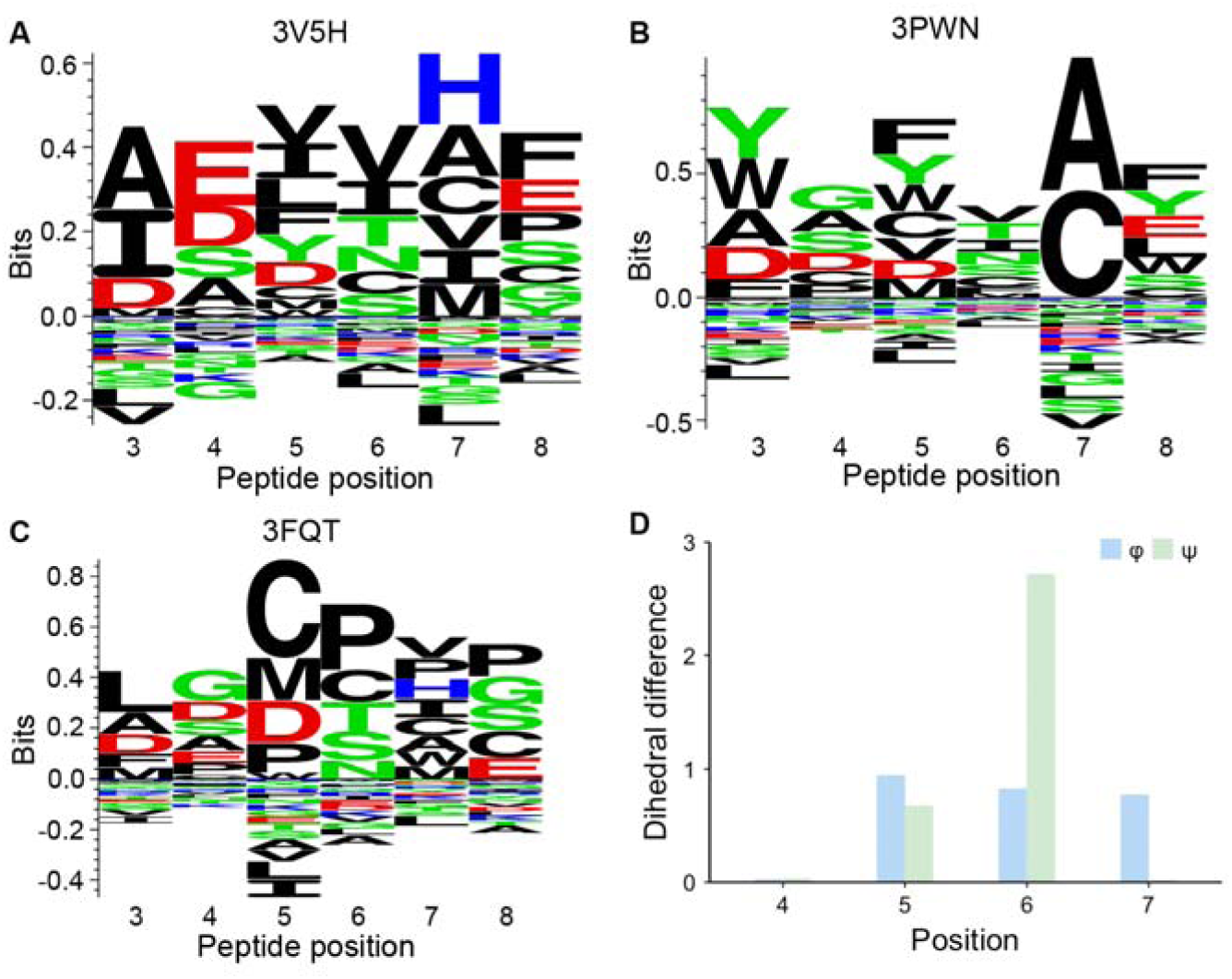
Additional analysis of the exhaustive structural modeling results. Peptide sequence logos from structural modeling results (n = 7,840) of **(A)** 3V5H, **(B)** 3PWN, and **(C)** 3FQT. Created using Seq2Logo. **(D)** Dihedral difference by position and angle for 5HHQ and 6VR1.

**Extended Data Figure 7.**
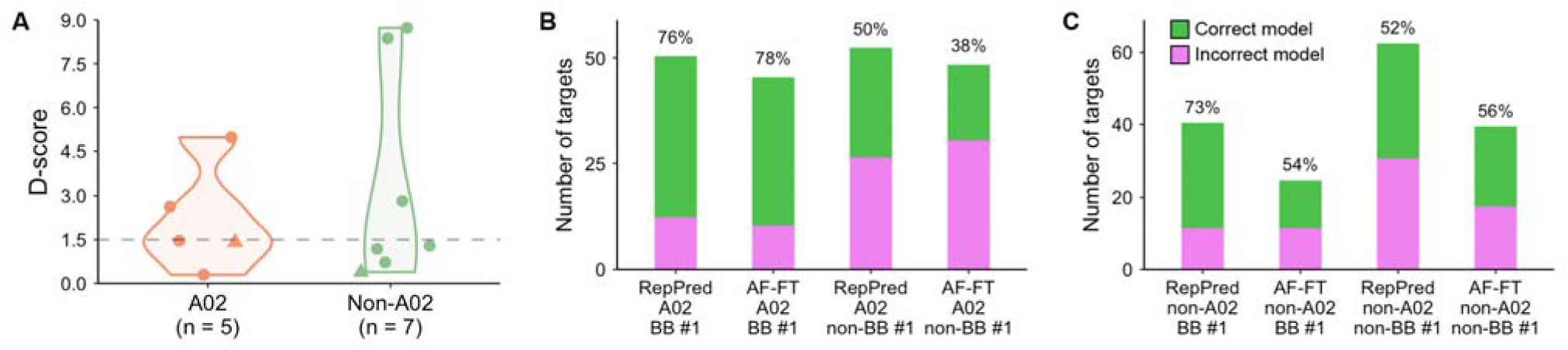
Additional analysis of the accuracy of RepPred and its comparison to AF-FT. **(A)** Overall results of blind testing of RepPred. Points depicted as triangles are elaborated in Fig. 4C and 4E. Comparison of RepPred and AF-FT for the most common backbone conformation (PDB ID 6J1V) and all other backbones for **(B)** A02 targets and **(C**) non-A02 targets.

