## Supplemental Figures and Tables for "HLA3DB: comprehensive annotation of peptide/HLA complexes enables blind structure prediction of T cell epitopes"

**This PDF file includes:**

Supplemental Figure 1
Supplementary Table 1

### Supplementary Information

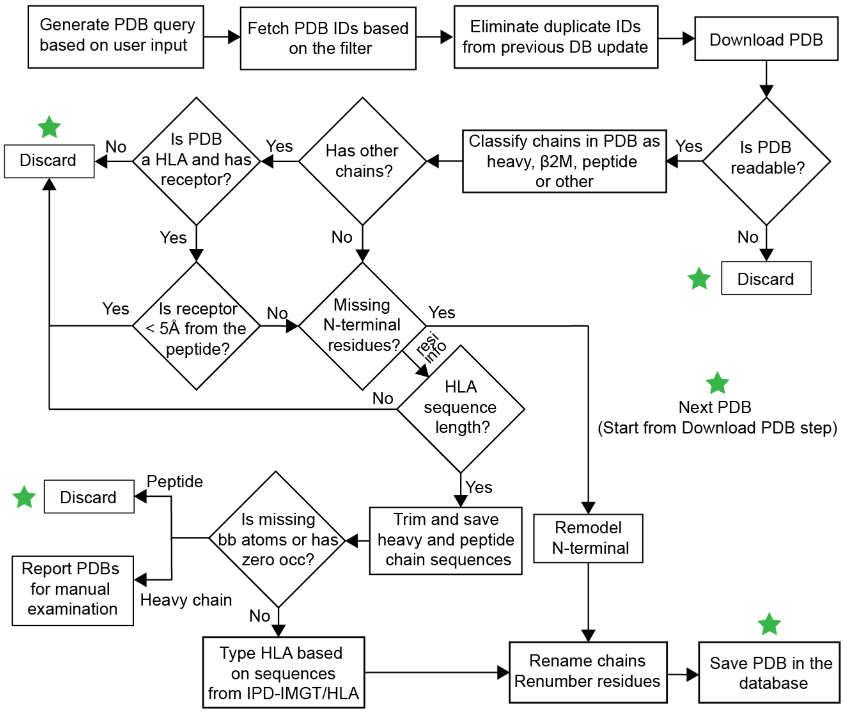


### Supplementary Figure 1

HLA3DB curation schematic implemented in Python using the RCSB PDB Search API.

| Peptide position | Amino acids |
|---|---|
| **3** | A, D, F, I, L, M, P, W, Y |
| **4** | A, C, D, E, G, P, S, W |
| **5** | C, D, F, I, L, M, P, V, W, Y |
| **6** | C, F, I, L, M, N, P, S, T, V, W |
| **7** | A, C, F, H, I, L, M, P, V, W, Y |
| **8** | C, E, F, G, L, P, S, W, Y |

**Supplementary Table 1**

Amino acids used for exhaustive structural modeling of likely peptide sequence combinations in the HLA-A*02:01 groove derived from peptides deposited in the IEDB.
